# Optogenetic control of the integrated stress response reveals proportional encoding and the stress memory landscape

**DOI:** 10.1101/2022.05.24.493309

**Authors:** Taivan Batjargal, Francesca Zappa, Ryan J Grant, Robert A Piscopio, Alex Chialastri, Siddharth S Dey, Diego Acosta-Alvear, Maxwell Z Wilson

## Abstract

The Integrated Stress Response (ISR) is a conserved signaling network that detects cellular damage and computes adaptive or terminal outcomes. Understanding the mechanisms that underly these computations has been difficult because natural stress inputs activate multiple parallel signaling pathways and classical ISR inducers have pleiotropic effects. To overcome this challenge, we engineered photo-switchable control over the ISR stress sensor kinase PKR (opto-PKR), which allows virtual control of the ISR. Using controlled light inputs to activate opto-PKR we traced information flow in the ISR both globally, in the transcriptome, and for key ISR effectors. Our analyses revealed a biphasic, input-proportional transcriptional response with two dynamic modes, transient and gradual, that correspond to adaptive and terminal ISR outcomes. Using this data, we constructed an ordinary differential equation (ODE) model of the ISR which predicted system hysteresis dependent on prior stress durations and that stress memory encoding may lead to resilience. Our results demonstrate that the input dynamics of the ISR encode information in stress levels, durations, and the timing between stress encounters.

## Introduction

The Integrated Stress Response (ISR) is a conserved signaling network that allows cells to detect and react to cell-intrinsic and environmental stresses including nutritional deficits, mitochondrial dysfunction, redox imbalances, viral infections, harmful RNA conformers, and protein-folding perturbations in the endoplasmic reticulum (ER). Because of its fundamental nature, it is not surprising that dysregulation of the ISR has been implicated in the pathogenesis of our most widespread diseases, including diabetes, cancer, and neurodegeneration (Costa-Mattioli and Walter, 2020). The mammalian ISR is governed by four stress sensor kinases—PERK, PKR, HRI, and GCN2—all of which actuate signaling by phosphorylating a single serine of the α subunit of the eukaryotic translation initiation factor eIF2 (eIF2α), a heterotrimeric GTPase required for translation initiation. Phosphorylated eIF2α (p-eIF2α) is a potent competitive inhibitor of its guanine nucleotide exchange factor eIF2B (Adomavicius et al., 2019; Kashiwagi et al., 2019; Krishnamoorthy et al., 2001) thereby leading to a global shutdown of protein synthesis. Concomitantly, phosphorylation of eIF2α promotes the translation of select mRNAs, including those encoding the ISR effector transcription factors ATF4, CHOP, as well as GADD34, the regulatory subunit of protein-phosphatase 1 that mediates negative feedback on the system through dephosphorylation of eIF2α to subdue ISR signals (Harding et al., 2000; Lee et al., 2009; Lu et al., 2004; Novoa et al., 2001; Palam et al., 2011; Vattem and Wek, 2004). While the molecular structure and circuitry of the ISR in health and disease have been extensively studied (Reviewed Costa-Mattioli and Walter, 2020), major questions remain about how the ISR encodes information at the systems level. How does the intensity and duration of a stress determine the cell’s response? On what timescales and with what subnetworks does the ISR determine cell fate? How do past stresses influence cellular resilience?

These questions are difficult to address with classical ISR-inducing agents such as chemicals (e.g., the ER calcium reuptake inhibitor thapsigargin (Thastrup et al., 1990) and sodium arsenite (Lowenstein et al., 1991)), viral infection or synthetic RNAs (e.g., poly-I:C (Tamassia et al., 2008)), and physical stimuli such as heat-shock, because all of these stressors introduce pleiotropic changes coupled to molecular damage that conflate *bona fide* changes that result from the cell’s response to stress with changes resulting from failing cellular machinery. In addition, the dose-response relationship between real stressors and pathway activation is non-trivial because of the cascading failures that non-linearly alter cellular stress levels. Last, damage induced by stressors is not immediately reversible. Instead, it must be repaired by the cell, constraining the investigation of how the ISR decodes dynamic inputs.

An ideal approach for dissecting the ISR would have three properties: It would (1) isolate a single stress sensor and activate it virtually to avoid conflation of inputs from parallel pathways and molecular damage, (2) enable high-resolution input control to query input-output proportionality, and (3) exert precise control of recurrent stimulation dynamics to interrogate the relationship between past stresses and future ones.

Here, we develop an approach with exactly these properties, using an optogenetically engineered ISR kinase to dynamically stimulate the ISR pathway with “virtual stress.” Optogenetic control enables the selective activation of a single kinase of the ISR pathway without inducing real molecular damage, greatly reducing the combinatorial complexity of the signals that are turned on by real stressors. By changing the intensity of light stimulation, it can tune the fraction of optogenetic proteins in the activated state, thereby precisely altering the perceived intensity. Light can also be quickly toggled on and off in a matter of milliseconds, enabling the delivery of precisely defined input dynamics. By assaying the global cellular changes in gene expression and the biochemical status at key protein nodes, such as eIF2α, in response to virtual stress of varying intensities and durations using high-throughput 96-well light delivery devices (Bugaj and Lim, 2019), we can use optogenetic control to dissect the complex computations that govern the ISR pathway.

We apply our approach to the ISR kinase PKR, an innate immunity effector of vertebrates that detects viral and endogenous double stranded RNA (Ben-Asouli et al., 2002; Chung et al., 2018; Kim et al., 2018; Thomis and Samuel, 1993; Youssef et al., 2015). PKR signaling is dysregulated in numerous diseases, including cancer and neurodegenerative disorders (Gal-Ben-Ari et al., 2018; Martinez et al., 2021), illustrating its central role in maintaining organismal homeostasis. Thus, the interrogation of stress encoding and decoding through optical control of PKR has both a broad significance and specific implications for understanding the fundamental mechanisms cells use to respond to stress.

While the role of ISR dynamics has not been systematically interrogated, observations suggest it utilizes the modalities of dose and dynamics to encode a wide variety of cell states. PKR-induced ISR activity has been shown to stochastically pulse on the timescales of hours in response to viral infections suggesting host-pathogen modulation of ISR dynamics (Ruggieri et al., 2012). More generally, the difference between an adaptive and a maladaptive stress response is commonly defined by the difference in dosage and timing of a stress input (Murray et al., 2022) (e.g. low, repeated doses of stress induce acclimation whereas larger chronic doses cause physiological dysfunction). Both examples suggest that hysteresis, history-dependent dynamics (Strogatz, 1994), in the ISR may play a functional role in dictating outcomes.

To understand how the ISR decodes stress signaling features we systematically altered both the levels and dynamics of opto-PKR activity. We coupled these precise input control methods with quantitative, time-resolved analysis of global transcriptomic changes as well as single-cell measurements of individual ISR effectors. This strategy revealed transcriptome remodeling governed by two dynamical modes–transient and gradual–that are associated with adaptive and terminal ISR outcomes. By systematically varying virtual stress intensity we determined that p-eIF2α, the adaptive transcription factor ATF4, and the pro-apoptotic transcription factor CHOP, all respond proportionally, but with fixed temporal phases. We leveraged these insights to develop an ordinary differential equation (ODE) model which quantitatively predicted that both the duration of past stress and the recovery time after stress determine cellular stress memory. Experimental validation of this model mapped the landscape of cellular adaptation to stress.

## Results

### Optogenetic activation of PKR induces the ISR

We chose to engineer optogenetic control over PKR for two reasons. First, PKR’s activation mechanism involves high-order self-association (Corbet et al., 2022; Mayo et al., 2019; Zappa et al., 2021), which can be mimicked using optogenetic clustering tools. Second, PKR is cytoplasmically localized and its sensor domain is isolated to a contiguous stretch of amino acids on the polypeptide chain, making it easier to engineer in comparison to the other ISR kinases (Cole, 2007; Zhu et al., 1997). Thus, we hypothesized that clustering PKR through exchanging PKR’s double-stranded RNA binding domains (dRBM1 and dRBM2) with photo-switchable variants of Cryptomchrome2 (Cry2), a light-inducible oligomerizer, would result in light-based activation of the ISR (Fig. 1A).

**Figure 1.**
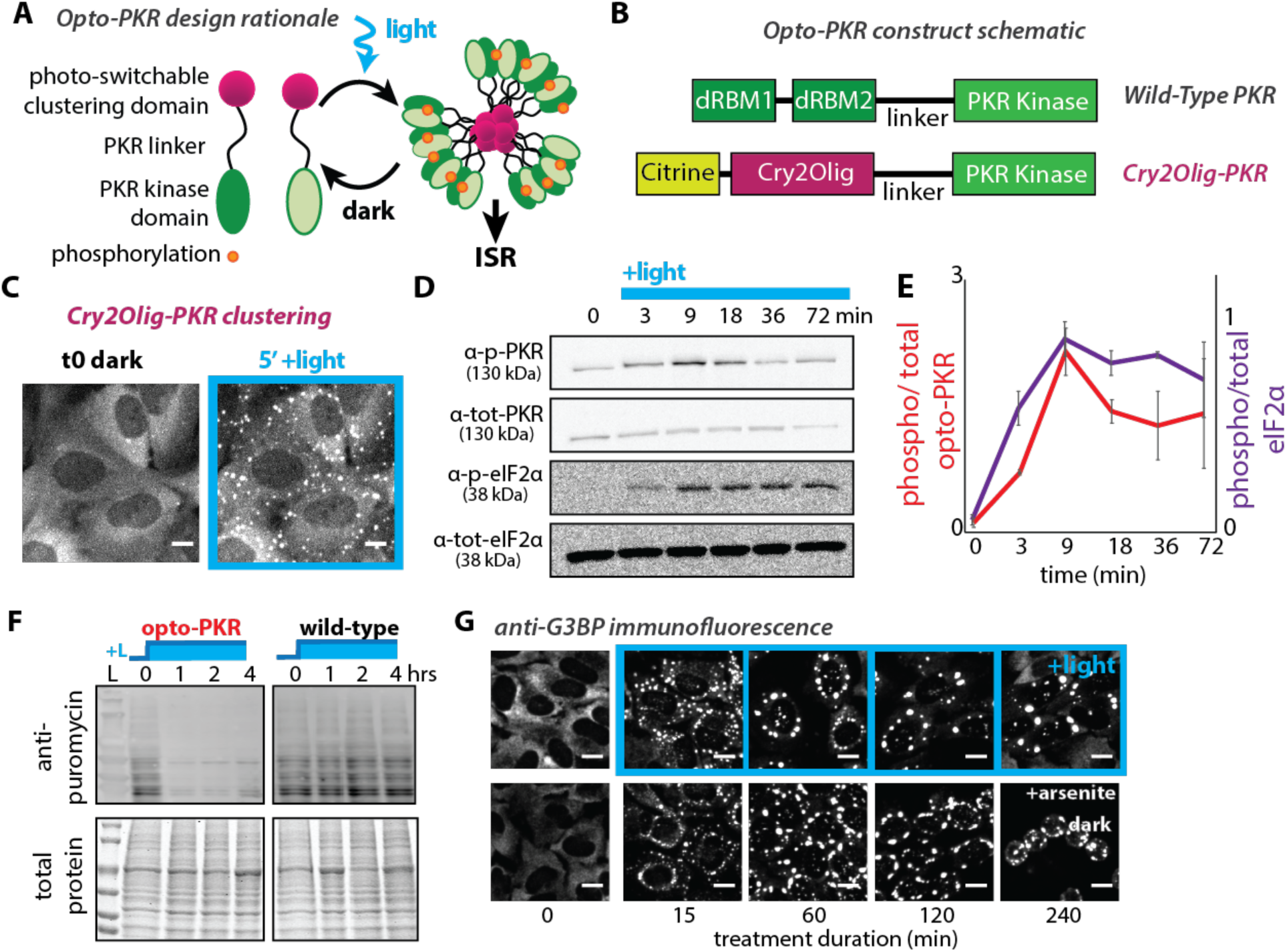
Optogenetic clustering of opto-PKR induces the canonical ISR. (A) Schematic representation of opto-PKR-driven induction of the ISR. The light-responsive Cry2 domain reversibly oligomerizes leading to clustering and trans-autophosphorylation of opto-PKR (blue light) or shutdown of the response (no light). (B) Schematic of the construct encoding opto-PKR. Cry2Olig domain fused to human PKR at residue 170 to 551. dRBM1/2, double-stranded RNA binding motifs. (C) Live cell imaging of opto-PKR in H4 cells under constant 448 nm illumination. Images are the maximum intensity projection of 5 Z-planes taken through the cell. Scale bar = 10 µm (D) Representative Western blot of phosphorylated and total opto-PKR detected using antibodies against total and phosphorylated PKR, and eIF2α under continuous blue light stimulation. (E) Quantification of the data in D. Phosphorylation signals were normalized as the ratio to the total protein levels (PKR or eIF2α) at each time point. N = 3. Error bars: SEM. (F) Western blot of puromycin incorporation into nascent peptides under blue light conditions in cells bearing opto-PKR compared to wild-type. Note the string translational repression elicited by active opto-PKR. L: light stimulation. (G) Immunofluorescence analysis of the SG marker G3BP1. Representative images of opto-PKR cells exposed to blue light or in the dark and treated with sodium arsenite (500 µM). Note that arsenite induces cell death while light stimulation does not. Scale bar = 10 µm.

To construct optogenetic PKR (opto-PKR) variants we replaced dRBM1 and dRBM2 with one of three optimized variants of Cry2: Cry2Olig, Cry2OClust, and Cry2ODrop (Fig. 1B, Fig. S1A). All three variants contain the E490G mutation in Cry2, which enhances the proteins’ oligomerization efficiency and is commonly referred to as Cry2Olig (Taslimi et al., 2014). In addition to this mutation, Cry2OClust and Cry2ODrop have C- and N-terminal peptide sequences, respectively, that increase their dynamic exchange with the cytoplasm (Park et al., 2017; Shin et al., 2017) (Fig. S1A). In this manner, Cry2Olig-PKR, Cry2OClust-PKR, and Cry2ODrop-PKR sample a range of biophysics, from more solid-like to more liquid-like condensates.

To test if our opto-PKR variants formed light-induced clusters we transduced H4 neuroglioma cells as well as U2OS osteosarcoma cells with each construct, sorted them for expression-matched populations using fluorescence-activated cell sorting (Fig. S1B) and took timelapse images while continuously stimulating them with activating light. All three opto-PKR variants in both cell lines formed condensates demonstrating light-responsive clustering of the engineered proteins (Fig. 1C, Fig. S1C,E, Supp Vid. 1). We noted that the number and size of condensates rapidly increased and then declined for all opto-PKR constructs in both cell lines (Fig. S1C-E). An established point mutation (K296R) in the phosphate transfer site of the kinase domain of PKR (Thomis and Samuel, 1993) (Fig. S1F) stabilized these clusters indicating that their activation-induced dissolution requires kinase activity (Fig. S1G, Supp Vid. 2).

Next, we sought to test whether our opto-PKR variants faithfully induce a canonical ISR despite the dynamic changes in cluster size and number. We first confirmed that none of the opto-PKR variants were constitutively activated in the dark and that illumination of WT H4 cells did not induce phosphorylation of either PKR or eIF2α (Fig. S1H-I). We then measured the phosphorylation kinetics of engineered PKR (p-PKR, which we resolved on Western blots by its higher molecular weight than endogenous at 130 kDa) and endogenous eIF2α (p-eIF2α) by Western blot in response to continuous illumination. While all opto-PKR variants rapidly induced high autophosphorylation (p-PKR), we noted that Cry2Olig-PKR maintained higher levels of phosphorylation for the duration of the 72-minute experiment (Fig 1D-E, Fig. S1J-K). Cry2OClust-PKR and Cry2ODrop-PKR demonstrated a lower dynamic range and more pulsatile activation kinetics, respectively, at both the level of p-opto-PKR and p-eIF2α (Fig. S1K). Given its more consistent activation kinetics we chose Cry2Olig-PKR from here on, and we refer to this variant as “opto-PKR” in the remainder of our experiments.

We next sought to understand if opto-PKR activation induced canonical ISR-driven translation repression (Dever et al., 1992) and formation of stress granules (SGs), ribonucleoprotein complexes formed in response to p-eIF2α (Buchan and Parker, 2009). To assess translational repression, we continuously activated opto-PKR and measured puromycin incorporation into nascent peptides using an anti-puromycin antibody (Schmidt et al., 2009). Opto-PKR activation drastically reduced translation initiation measured by nascent peptide puromycilation (Fig. 1F), which is consistent with translational repression observed by ISR activation with chemical inducers (Dey et al., 2010). Next, we monitored SG formation by immunofluorescence (IF) of the SG assembly factor G3BP1 (Kedersha et al., 2016) upon activation of opto-PKR. Light-induced SGs were indistinguishable from those formed through arsenite induction, except for those observed at late time points (>4 hours) where arsenite treated cells began to detach from the plate and die (Fig. 1G, Fig. S1M). Altogether our data demonstrate that opto-PKR recapitulates the major ISR events across multiple cell lines, allowing us to virtually control the ISR.

### The stress-free ISR has a biphasic transcriptional response

Chemical ISR inducers have pleiotropic effects, rendering them ineffective tools to dissect the cellular responses to stress from cellular damage. To overcome this limitation, we deployed opto-PKR to globally characterize the dynamics of transcriptome remodeling in response to a “stress-free” ISR. We wondered whether distinct dynamic ISR phases might exist in response to virtual stress. To this end, we performed a time-resolved RNAseq experiment to characterize the dynamic changes in the transcriptome upon continuous stimulation with virtual stress. We utilized the optoPlate light delivery device (Bugaj and Lim, 2019), which allows for arbitrary control over the illumination sequence of each well of a 96-well plate, to vary the duration of 450 nm light (∼7.5 millicandela, see methods for voltages) (Fig. 2A). To obtain time-resolved transcriptional changes we sampled cells at 7 time points, from 0 to 12 hours.

**Figure 2.**
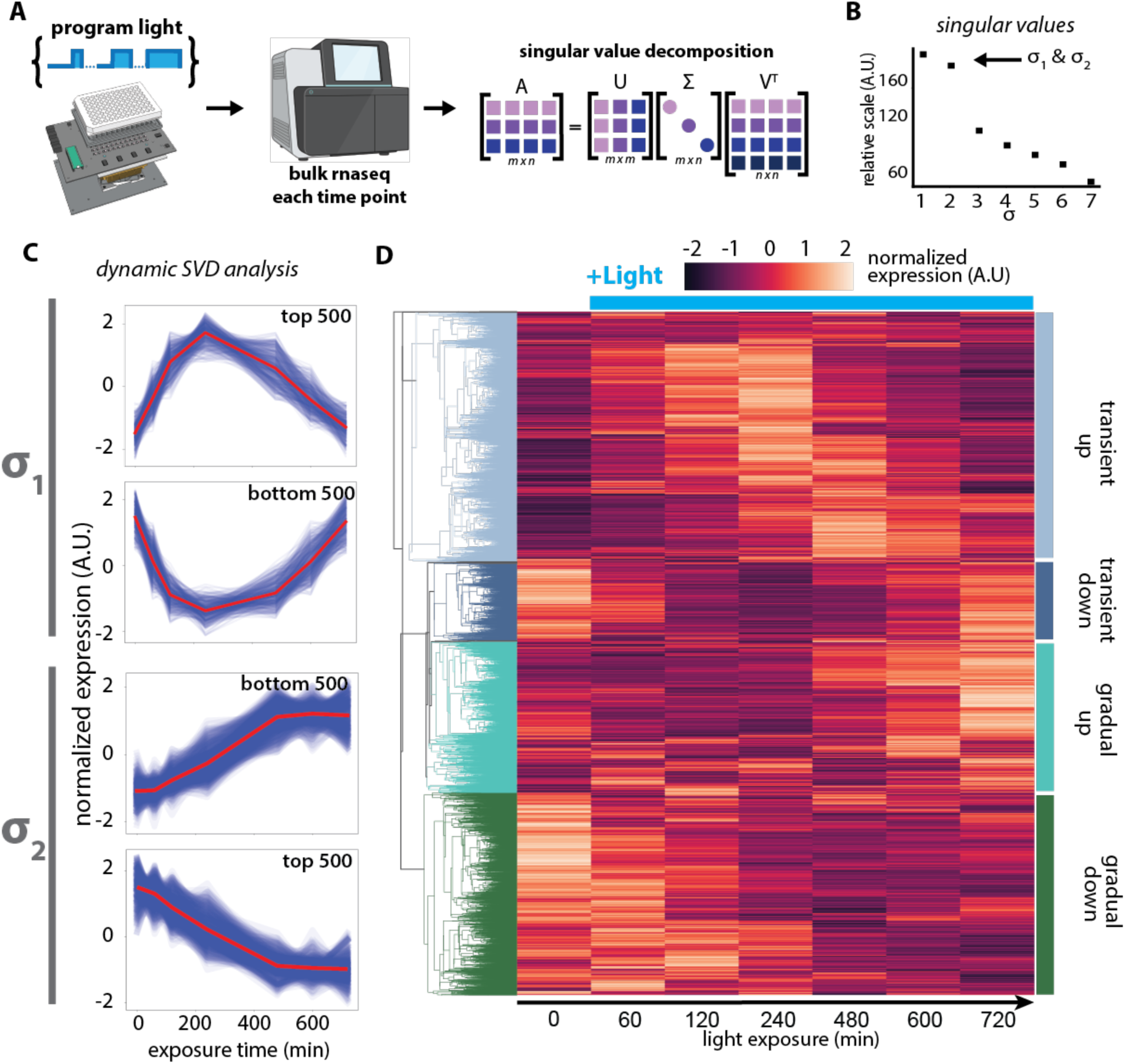
The ISR has a biphasic transcriptional response to virtual stress. (A) Experimental design. To investigate transcriptomic dynamics of opto-PKR-induced ISR over time, H4-cells bearing opto-PKR were illuminated using the optoPlate device to collect 7 samples across 720 mins. (B) Plot of singular values from Σ matrix of singular value decomposition on a relative scale(C) Plot of the top 500 and bottom 500 genes from each U vector associated with the most prominent two dynamic modes, σ_1_and σ_2_. Each blue line represents a distinct gene from our z-scored RNA-seq data, the red line is the mean of 500 plotted genes over time. (D) Centroid-clustered heatmap of the global transcriptome dynamics over 12 hours. Rooted tree diagram of gene clustering is color coded to indicate 4 dynamic modes.

Examination of individual markers of the ISR (*e*.*g. ATF3, DDIT3* (CHOP), *PPP1R15A* (GADD34), etc.) showed two distinct classes of dynamical responses corresponding with transiently pulsed and gradually accumulating mRNA kinetics (Fig. S2A).

These two classes of mRNA kinetics suggested that the global transcriptome dynamics may also display a similar pattern. Thus, to map the dynamical gene expression modes of the ISR we performed a Singular Value Decomposition on the Z-scored, time-resolved transcriptome matrix (Fig. 2A). We found that the first two modes (σ_1_ and σ_2_) explained more than half of the total variance of the transcriptome (Fig. 2B). To visualize these two dominant modes, we plotted the top 500 and bottom 500 genes from the U vectors associated with σ_1_ and σ_2_. This analysis showed that σ_1_ represented a transient response, with genes reaching their maximum between 4-6 hours of continuous activation (Fig. 2C *top*). On the other hand, genes represented by σ_2_ showed a gradual, but continuous change that leveled off between 8-10 hours of illumination (Fig. 2C *bottom*). Analyzing the transcriptome separately through centroid clustering recapitulated these two dynamic modes (Fig. 2D).

Gene ontology (GO) enrichment analysis of these modes hinted that transiently downregulated processes included translation, purine and carbohydrate metabolism, and gonadal mesoderm development, while transiently upregulated processes included protein ubiquitination and TGF signaling.

This analysis suggest that the transient mode is associated with adaptive remodeling of the proteome and alteration of cell identity. On the other hand, gradually upregulated genes were associated with apoptosis and ER and nutrient stress responses, while gradually downregulated processes included mitosis and cell adhesion (Fig. S2B). This analysis suggest that the gradual mode is associated with a switch towards a terminal response. The regulation of developmental signaling and adhesion by the ISR in neuroglioma cells may reflect oncogenic rewiring. Together, these results indicate that the ISR has two prominent dynamic modes–transient and gradual–and that these modes represent the adaptive and terminal responses of the ISR.

### Key ISR transcription factors respond proportionally with fixed phases

The ISR culminates in gene expression programs controlled by adaptive (ATF4) and pro-apoptotic (CHOP) transcription factors, yet the question of how these transcription factors respond to varying the magnitude of a stress has remained unanswered. This question has been difficult to probe with chemical stressors because they induce cascading failures that can lead to a non-linear relationship between chemical stressor concentration and ISR activity (Pakos-Zebrucka et al., 2016). The ISR has been characterized to be switch-like in some instances (Vattem and Wek, 2004), while recent findings regarding the mechanism of action of ISR-inhibiting drugs (e.g. ISRIB) suggest a more nuanced dose-response relationship (Costa-Mattioli and Walter, 2020; Tsai et al., 2018; Zyryanova et al., 2018). Thus, we wondered how the ISR signaling dynamics are affected by varying magnitudes of virtual stress inputs.

To this end, we characterized the dynamics, at the protein level, of p-eIF2α, ATF4, and CHOP, which represent the ISR core, and its adaptive and terminal phases, respectively (Fig. 3A). To control the magnitude of virtual stress we varied light intensity as it varies the proportion of Cry2 molecules in the photo-active state (Shin et al., 2017). Using the optoPlate device we applied light ranging from 0.2 to 7.5 mcd for over 10 hours (Fig. S3A) and collected time points for IF analysis of p-eIF2α, ATF4, and CHOP to obtain their endogenous dynamic responses (Fig. 3B). To analyze the IF images, we developed a custom image analysis pipeline that extracts single-cell fluorescence intensities from the nucleus, cytoplasm, and SGs, which we used to quantify the response in over 100 cells per condition for a total of more than 10,000 single cells across all times and intensities (Fig. S3B-C).

**Figure 3.**
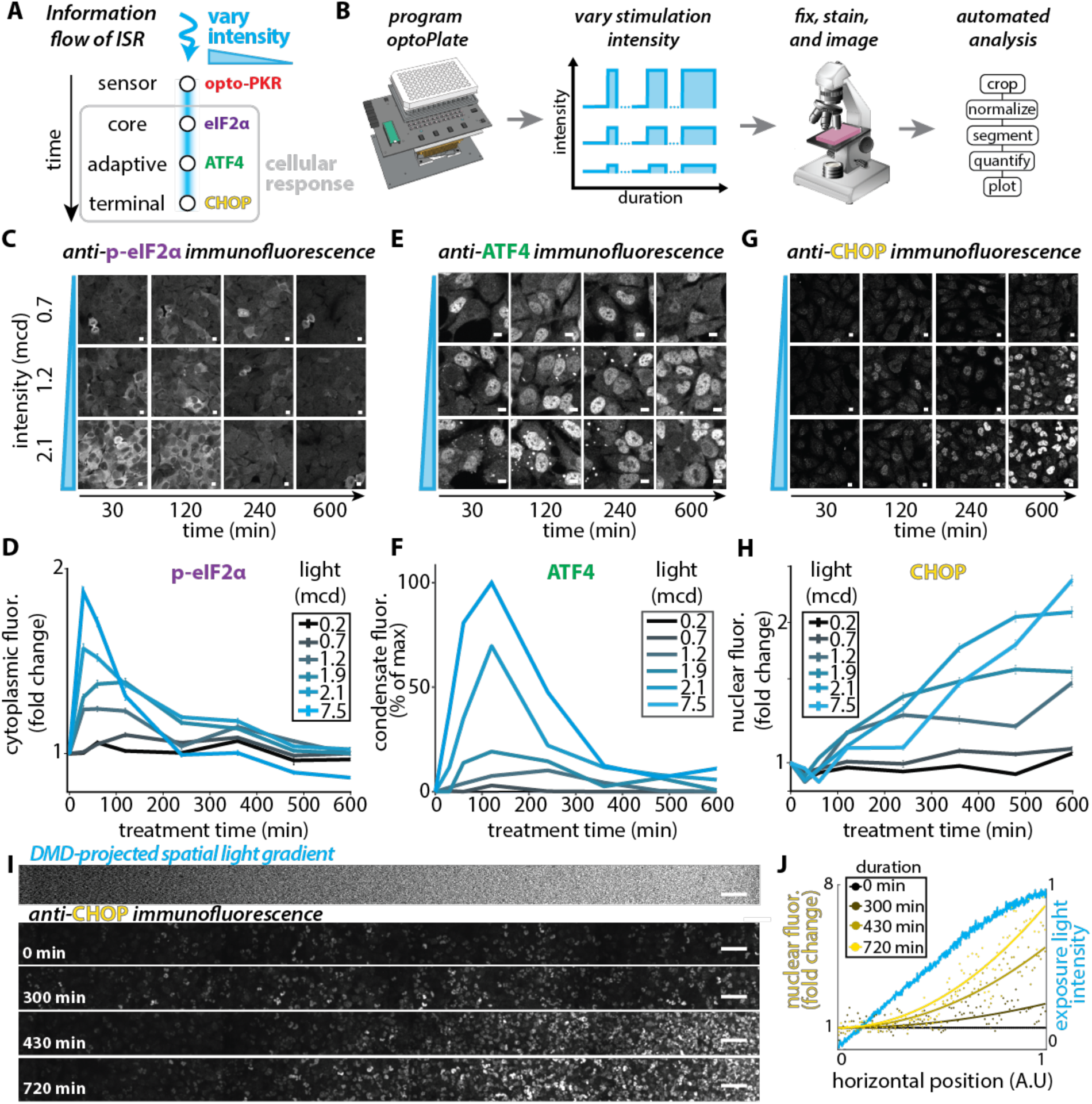
ISR transcription factors in different ISR signaling modes respond proportionally with fixed phases. (A) Schematic of ISR information flow through activation of opto-PKR. (B) Experimental design to query responses to varying illumination intensity over time. Blue light intensity was varied from 0.2-7.5 mcd using the optoPlate device over the period of 600 mins. Cells were fixed and immunostained for the indicated markers, imaged and immunofluorescence data was processed with an automated pipeline. (C,E,G) Representative immunofluorescence images of H4 cells bearing opto-PKR under the light exposures outlined, varying combinations of intensity and duration, stained for p-eIF2α (C), ATF4 (E) and CHOP (G), respectively. n > 150. Scale bar = 10 µm. (D,F,H) Quantification of immunofluorescence data in C,E,G. Each line corresponds to a specific illumination intensity over time. Error bars: SEM (I) Representative image of light-based gradient activation of opto-PKR in H4 cells immunostained for CHOP. The light gradient was applied using a digital micro-mirror device (DMD) and continuous illumination. Scale bar = 100 µm. (J) Quantification of nuclear CHOP signal upon gradient light stimulation. Each point corresponds to the mean nuclear immunofluorescence intensity of all nuclei positioned within one of hundred equally divided horizontal space regions. Each line is a best fit adhering to a quadratic formula: f(x) = ax^2+1. n > 2900.

In general, we found that the ISR nodes we examined had features that were modulated in proportion to their inputs, albeit with fixed phases. P-eIF2α peaked within 30 min of activation and then was repressed by 4 hours with the peak magnitude varying in proportion to light intensity (Fig. 3C-D). We found that ATF4 localized to SGs (Fig. S3B) as has been reported previously (Mateju et al., 2020). SG-localized ATF4 peaked at 2 hours of activation, where a subset of the adaptive response genes peak (Fig. S2A), with its peak height also proportional to the input (Fig. 3E-F). We also noted a proportional response in the change in nuclear ATF4, although its magnitude was much lower than SG-localized ATF4 (Fig. S3D). Finally, we found that CHOP accumulated gradually with its rate of accumulation varying with the input intensity (Fig. 3G-H).

We further probed the proportional intensity encoding of CHOP accumulation because of its pro-apoptotic role (Oyadomari and Mori, 2004). We applied a smooth (256 bit) linear gradient of illumination across a field of opto-PKR-bearing cells and IF-stained for CHOP at four time points ranging from 0 to 12 hours (Fig. 3I *top*). Remarkably, this stimulation led to a spatial gradient of CHOP (Fig. 3I). Quantification of the data showed a best fit corresponding with a 2^nd^ order polynomial, suggesting that the ISR takes the integral of stress inputs when computing CHOP levels (Fig. 3J, Fig. S3E). Overall, these experiments demonstrate that the peaks of p-eIF2α and ATF4 as well as the slope of CHOP are modulated in proportion to their inputs, while the timing of these responses are independent of input magnitude.

### A simple ODE model of the ISR captures core responses and predicts stress memory

Our studies so far have focused on probing the cellular response to continuous stress inputs. However, the ISR naturally dynamically oscillates (Ruggieri et al., 2012) and the proposed adaptive effects of stress are inherently multi-phasic, in that they describe the effect of past stresses on future ones (Calabrese and Calabrese, 2022; Calabrese et al., 2007; Ost et al., 2016). To predict how dynamic stress inputs are processed, we constructed and fit a simple ODE model of the ISR. While the ISR has been modeled before (Klein et al., 2022), we sought to build the simplest model that still captured the system dynamics and to be the first to parameterize a model using non-damaging inputs, which could confound the interpretation of the ISR’s dynamics.

We chose to represent the ISR as three interconnected modules: sensor, core, and memory (Fig. 4A). We approximate the sensor module, our ability to activate it with light, and its phosphorylation of eIF2α using mass action kinetics (Fig. S4A). To minimize the number of free parameters we simplified the core and memory modules by choosing to represent the GDP and GTP-bound forms of eIF2α, as well as the multiple eIF2B assembly states, as non-linear cooperative interactions between eIF2α, p-eIF2α, and the well-known phosphatase coupling protein GADD34 (Fig. 4A, Fig. S4A). To parameterize our model, we fit it to the normalized p-eIF2α data collected for the maximal continuous activation shown in Fig. 3D. We found that the fit, in terms of reduced Chi-Squared statistic, of our simple cooperative model was better than a model lacking the non-linear terms (Fig. S4B-C,E-F), justifying their inclusion. The model reproduced the negative feedback that drives p-eIF2α back to baseline in response to a continuous step input (Fig. 4B) as well as the proportional modulation of peak p-eIF2α without shifting its phase (Fig. 4C).

**Figure 4.**
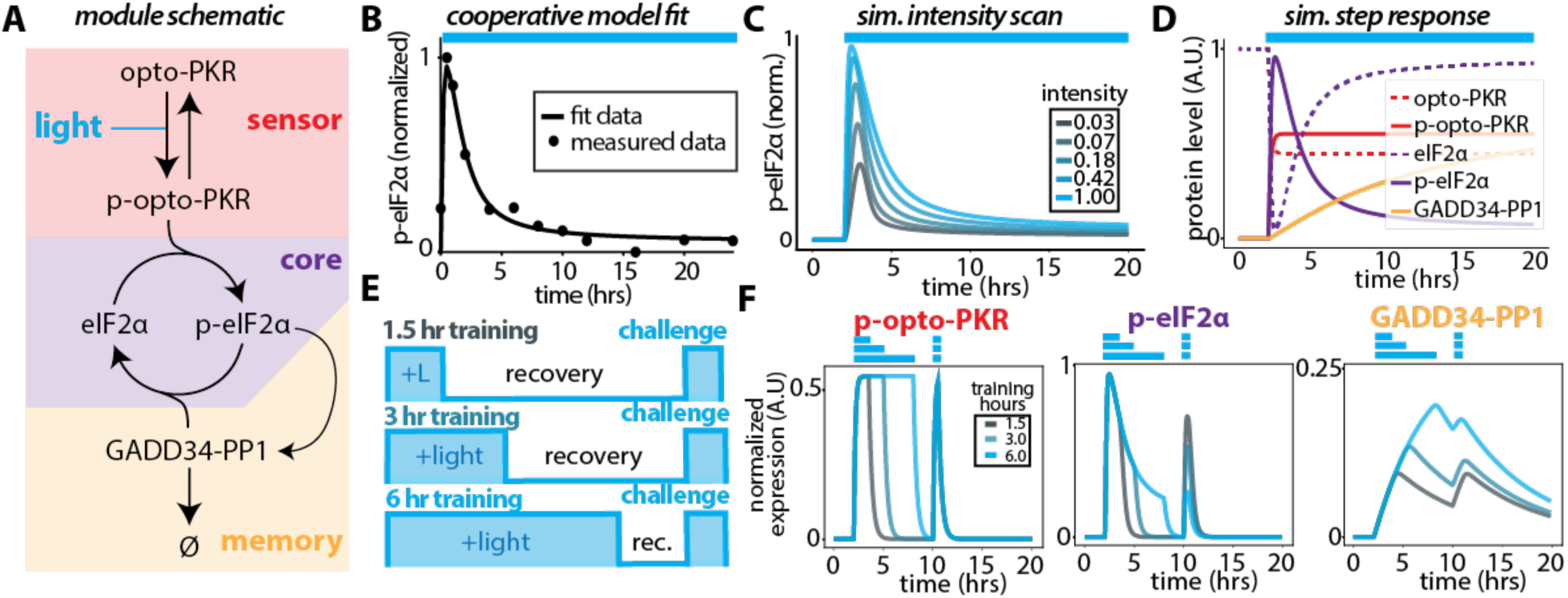
A simple ODE model of the ISR captures core responses and predicts stress memory. (A) Schematic representation of interactions simulated in our model using equations representing opto-PKR, p-opto-PKR, eIF2α, p-eIF2α and GADD3-PP1 complex, segregated to three sub-modules of the ISR. (B) Best fit of simulated data to the cooperative ODE model to experimental data for p-eIF2α. The line is simulated data, and the points are experimentally measured data. (C) Simulations showing changes in p-eIF2α levels in response to decreasing illumination intensity. (D) Simulated response of all state variables of our cooperative model after fitting to experimental data. (E) Demonstration of inputs used to probe system hysteresis. To investigate the effect of past stress inputs to present stress inputs, we simulated multiple increasing ratios of stress input to recovery in the same period, before applying an identical challenge to all simulations. (F) Simulated stress memory through modulation of input and recovery ratios. Normalized levels of key state variables over time: p-opto-PKR (left), p-eIF2α (mid), GADD34-PP1 (right). Challenge occurs at 10 hours for all simulations.

Examining the other state variables in response to continuous activation we found that the GADD34-PP1 complex accumulated over the course of 20 hrs (Fig. 4D). We hypothesized that this background accumulation of effective PP1 activity could desensitize future inputs as a function of past inputs. To test this hypothesis, we simulated a short (1.5 hrs), intermediate (3 hrs), and long (6 hrs) stimulation followed by a uniform 30 min challenge, corresponding to the peak activation of the naïve ISR (Fig. 4E). These simulations demonstrated a negative relationship between the amount of prior stimulation on the ability of the core module of the ISR to respond to future challenges, despite the sensor module responding identically for all inputs, which was mediated by the varying accumulation of GADD34-PP1 in the memory module (Fig. 4F). Overall, our model recapitulates the observed ISR dynamics and suggests that the ISR can encode memories of prior stimulations in the GADD34-PP1 feedback control module as has been recently suggested (Klein et al., 2022).

### Input dynamics shape the stress memory landscape

Chronically stressed cells respond differently than those exposed to acute stress (Guan et al., 2017), suggesting a role for stress input duration in shaping the ISR. Our simulations predicted prior inputs attenuate the system’s response to future inputs, thus constituting a stress memory. To map the stress memory landscape, we varied the duration of stress input independently from the recovery duration. This stimulation/recovery regime was followed by a uniform challenge (10 min input) to query the cells’ response (Fig. 5A). Using our model, we applied these variable training and recovery times to simulate the stress memory landscape. We found that the p-eIF2α response was dependent on both conditioning variables (Fig. 5B) and anticorrelated with GADD34-PP1 levels at the time of challenge (Fig. S5C), as expected. Notably, the response of the ISR to the fixed challenge in simulations that were trained with long durations of stress (∼3-10 hrs) but given short recovery times (< 7 hrs) was severely blunted (Fig. 5B).

**Figure 5.**
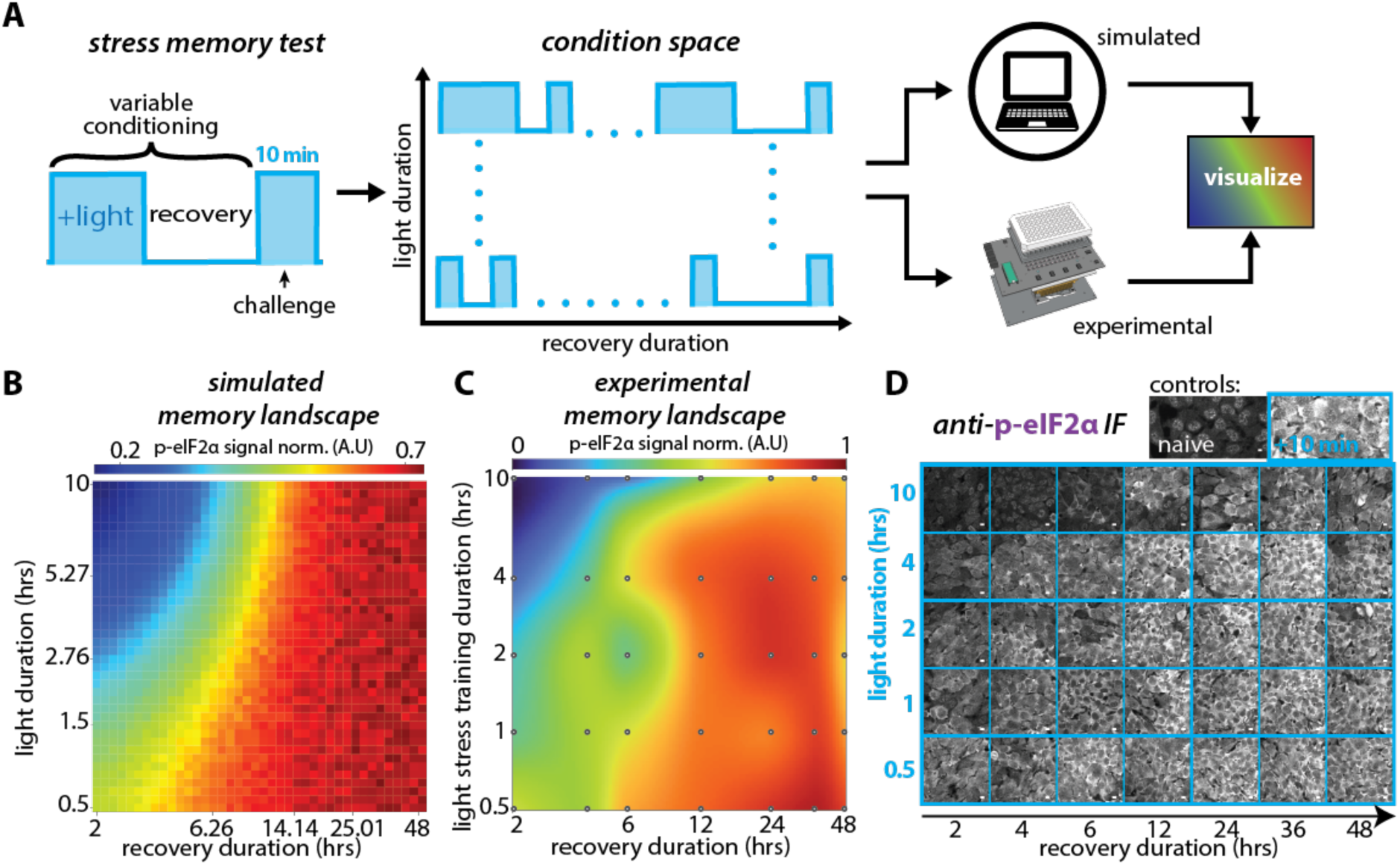
Input dynamics shape the stress memory landscape. (A) Experimental scheme used to map the stress memory landscape. Cells were conditioned by varying stress input and recovery durations, before applying an identical challenge to query present response. Conditioning inputs were applied to the cooperative model and to cells for comparison. Cells were illuminated with the optoPlate device, fixed, immunostained, and quantified for p-eIF2α (similarly to 3B). (B) Heatmap of simulated stress memory landscape for p-eIF2α in response to 10 min challenge. Total duration of conditioning varies. (C) Interpolated heatmap of experimental stress memory landscape for p-eIF2α in response to 10 min challenge. Experimental data quantified as normalized mean intensity of cytoplasmic immunofluorescence signal. Each dot represents a sample point. n > 375 single cells for each point. (D) Representative immunofluorescence images immunostained for p-eIF2α after conditioning and challenge. Controls: naïve = no illumination, +10 min = only challenge (no conditioning). Scale bar = 10 µm.

To test if our model predicted the stress memory landscape, we used the optoPlate device to probe the response to a subset of the conditioning inputs, followed by the same 10 min challenge. These experiments require rapid reversibility of the input to ensure that the queried cellular state is only a response to the most recent challenge and not to still decaying training inputs. Therefore, we first measured the decay of p-opto-PKR in response to a 20-minute pulse of activating light and found that it decayed on the timescale of 3 mins, an order of magnitude shorter than the briefest recovery duration tested (2 hrs) (Fig. S5A-B). We found that cells responded in accordance with our model prediction with long duration inputs and short recovery times severely blunting p-eIF2α levels (Fig. 5C). Short duration inputs also reduced p-eIF2α levels whereas cells were generally fully recovered after > 12 hrs (Fig. 5C). Upon examination of IF images, we found these patterns were visibly evident (Fig. 5D). Finally, we co-stained for the SG marker G3BP1, and found that only after the longest-duration inputs, corresponding to peak terminal phase, did cells completely inhibit SG formation (Fig. S5D). Together, our simulations and experimental data describe the timescales of cellular adaptation to stress and implicate GADD34 kinetics in determining the memory landscape.

## Discussion

Neuronal optogenetic tools revolutionized neuroscience because of their ability to causally dissect the role of neural firing dynamics in driving a brain or behavior state. Here we use cellular optogenetic tools analogously, to control the perceived ISR input and understand how ISR signaling dynamics dictate cellular outcomes. With the ability to deliver arbitrary ISR input dynamics without damaging cells—we call this virtual stress—we were able to address three longstanding questions: (i) What are the temporal phases of the ISR? (ii) How do these phases alter the ISR output with varying stress intensities? And (iii) How is the ISR modified by its history?

Based on our analysis of time-resolved transcriptomic profiling, we propose that the ISR can be delineated into two distinct dynamical modes: (1) a transient mode associated with the pro-adaptive response that peaks at ∼4 hours and (2) a gradual mode that slowly activates terminal response genes in the background, leveling off at ∼10 hours. Using this insight, we probed pathway encoding at key nodes, representative of the gradual and transient phases, by coupling a high-throughput light delivery device to custom automated image analysis algorithms. This approach allowed us to systematically scan through activation levels and durations in thousands of single cells revealing that they respond proportionally to stress levels, albeit with their own phase-dependent dynamics. Surprisingly, we found that the timing of these two phases was unchanged by intensity, suggesting that stress duration is a central feature sensed by the ISR.

This interpretation of our data guided us to construct a simple ODE model of the ISR to explore its dynamical properties. We found that the model displays hysteresis, which suggested that it could be used to predict the encoding of stress memories. By systematically modeling variations in the duration of both stress input as well as recovery time, we simulated a stress memory landscape that predicted that longer stresses and shorter recovery times generally blunt the cell’s present response. We validated our model’s prediction experimentally, generating the first map of the stress memory landscape. Rather than focusing solely on cellular contexts where stress is delivered in a sustained manner using small molecule poisons, our work points to the importance of pulsatile and/or stochastic ISR dynamics that have now been observed to occur in response to viral infection (Klein et al., 2022).

### The value of ‘virtual’ stress in interrogating pathways that detect molecular damage

Virtually activating the ISR allowed us to systematically screen through fixed intensities of perceived stress making way for the analysis of a “stress-free” ISR. This allowed us to examine the global transcriptomic state of cells without confounding alterations due to failing cellular subroutines (such as a damaged ER resulting from thapsigargin treatment (Thastrup et al., 1990), or sublethal oxidation arising from arsenite exposure (Lowenstein et al., 1991)). By isolating a single sensor, PKR, our approach also allows us to eliminate any possible contributions of parallel stress response pathways that are activated by the same input, as occurs with multiple innate immunity dsRNA sensors (Hur, 2019). Interestingly, despite continuously activating the ISR for over 16 hours we never observed significant signs of cell death in the cell lines we studied. This could be explained by either that cancer cells have (i) re-wired their stress response to avoid or actively suppress apoptosis, (ii) that in certain cell types ISR-induced programmed cell death requires additional inputs which are satisfied by true damage, or (iii) that cell-death requires longer chronic ISR induction than was tested in this study. Indeed, GO analysis of the terminal phase showed apoptotic processes (GO:0006915) were significantly enriched in the upregulated gene class, indicating a potential preamble to ISR-induced apoptosis. Further, studies of how the ISR is re-wired across a range of cancers using orthogonally-controlled stress sensors such as ours could reveal how oncogenic transformation selects for genotypic and epigenetic states that curb ISR-induced cell death.

An additional advantage of optogenetic induction over traditional stressors is that it can be patterned arbitrarily in space through the spatial modulation of light. We used this property to probe the decoding function of CHOP. Using digital micromirror devices with 256-bit resolution we delivered a linear gradient of light to smoothly modulate the ratio of opto-PKR molecules in the photo-active and photo-inactive states. Cells produced a 2^nd^ order quadratic response in their CHOP levels, suggesting that cells take the integral of the stress experienced to compute CHOP outputs.

### The stress memory landscape in terms of normal cellular functions, adaptation, and stress related diseases

Our model and experiments not only make quantitative predictions but also suggest where stress memory is stored and what parameters should be modified to alter the shape of the stress memory landscape. Our simulations demonstrate that the instantaneous level of the GADD34-PP1 complex is, at least in part, responsible for the ISR’s hysteresis (Fig. 4E-F). However, at least another PP1 regulatory subunit CreP also induces dephosphorylation of eIF2α outside the negative feedback loop established by induction of GADD34 (Jousse et al., 2003), and should be accounted for in further evaluation of stress memories. We found that a cooperative term describing the rate of accumulation of the GADD34-PP1 complex fit the dynamics of p-eIF2α better than a simple mass action term (Fig S4C). This suggests that altering the four parameters of this cooperativity function (K, K_1_, K_2_, and ß_1_) as well as the degradation rate of GADD34 (ß_2_) will enable manipulation of the memory landscape. While the rates of accumulation and decay of GADD34 explain, to some degree, the encoding of stress memories (this work and Klein et al, 2022), we cannot discard the possibility that other regulators of the availability of ternary complex (i.e., CreP, eIF2B) could also contribute to the stress memory landscape. Drugs targeting these parameters, for example ISRIB, may prove to be useful in alleviating disease symptoms caused by failure of the cell to either forget or remember past stresses. Further *in vivo* studies will be required to address this possibility.

In healthy cells challenged with short durations of stress the ISR induces an adaptive response that increases resilience. On the other hand, this same pathway, when chronically activated causes disease marked by reduced cellular function and increased cell death (Martinez et al., 2021). Since the severity and likelihood of onset of many ISR-related diseases (e.g., Alzheimer’s, Huntington’s, Parkinson’s, and diabetes) increases with age, this observation begs the question of whether the ISR’s dynamical information processing capability is prone to corruption during aging. Recently, the role of altered signaling dynamics was shown to explain how some cancers misperceive extracellular signals, transmuting healthy dynamical information into oncogenic signals (Bugaj et al., 2018). Whether altered ISR dynamics explain disease states in a similar manner, with age-related changes in the ISR network altering the healthy cellular perception of stress, remains an open question. Future studies combining precise optogenetic stimuli with live-cell fluorescent reporters of canonical stress responding nodes, such as fluorescent protein CRISPR-tagged ATF4 and CHOP, in models of ISR-associated diseases may illuminate this question.

## Methods

### Cell culture

Human osteosarcoma epithelial cells (U2OS) and H4 neuroglioma cells (H4) were cultured in Dulbecco’s Modified Eagle Medium (DMEM) containing 4.5 g/mL glucose (Gibco), 10% fetal bovine serum (Atlas Biologicals, F-0500-D), 1% penicillin and streptomycin at 37°C and 5% carbon dioxide. All cells carrying light responsive proteins were maintained and treated in the absence of blue light (dark or red light only). Cell culture vessels containing light responsive cells were wrapped in aluminum foil to prevent accidental light leakage. Prior to microscopy experiments, cells were plated onto glass bottom dishes pre-treated with bovine fibronectin to allow for adhesion.

### DNA constructs

Plasmids were obtained from Maxwell Z. Wilson and Diego Acosta-Alvear. The DNA regions of interest were amplified from host plasmids by PCR using CloneAmp HiFi PCR Premix (Takara, 639298) with custom primers for Gibson assembly (with overhangs). Constructs of interest were generated through subcloning into a lentiviral expression vector bearing the SFFV promoter using Gibson Assembly as per the manufacturer’s recommendations. The resulting lentiviral expression vectors contained the chimeric PKR constructs. Constructs are available on Addgene.

### Lentivirus transduction

Lentiviral supernatant was produced by transfecting second generation helper constructs, pCMVdR8.91 and pMD2.G (gift of Jared Toettcher, Princeton University), together with our lentiviral constructs at a mass ratio of 0.444 : 0.056 : 0.500, respectively, into HEK-293T cells. Transfections were carried out using polyethyleneimine. HEK-293T cells were maintained for 48 hours post transfection at which point the supernatant was harvested, filtered through 0.45 µm filter and stored at -80 prior to infection of target cells.

### Fluorescence activated cell sorting

Engineered cells were sorted using Sony SH-800 cell sorter system. Gating was determined using untransduced parental cells as a negative control, and selected cells were sorted through a narrow fluorescence gate to obtain a pseudo-clonal cell line. All engineered cell lines used in this study originated from expansion of pseudo-clonal cell lines.

### Western blotting

Roughly 1 × 10^6^ cells per experimental condition were harvested using 1x Laemmli buffer (30 mM Tris-HCl pH 6.8, 1%SDS, 10% (w/v) Glycerol, bromophenol blue) from a single well of a 6-well plate immediately after treatment. The lysates were heated at 95°C for 3 minutes, briefly sonicated, supplemented with 5% 2-mercaptoethanol, and stored at -20°C conditions until SDS-PAGE. Samples were separated in 10% SDS-PAGE gels run at 100V for 1-2 hours, transferred onto PVDF membranes using a TE 77 semi-dry transfer unit (Cytiva Amersham) at 10V for 15 min. The membranes were blocked with 3% BSA in TBS-T for 1 hour, followed by an overnight incubation with primary antibody at 4°C. Membranes were washed and incubated with fluorescently or HRP conjugated secondary antibodies for 1 hour at room temperature. All incubation steps were followed by 3x washes with TBS-T for 10 min.

The membranes were imaged using enhanced chemiluminescence using a Radiance Plus imager (Azure Biosystems) or a Li-Cor Odyssey instrument. All antibodies were diluted in 3% BSA in TBS-T.

List of antibodies used in western blotting:

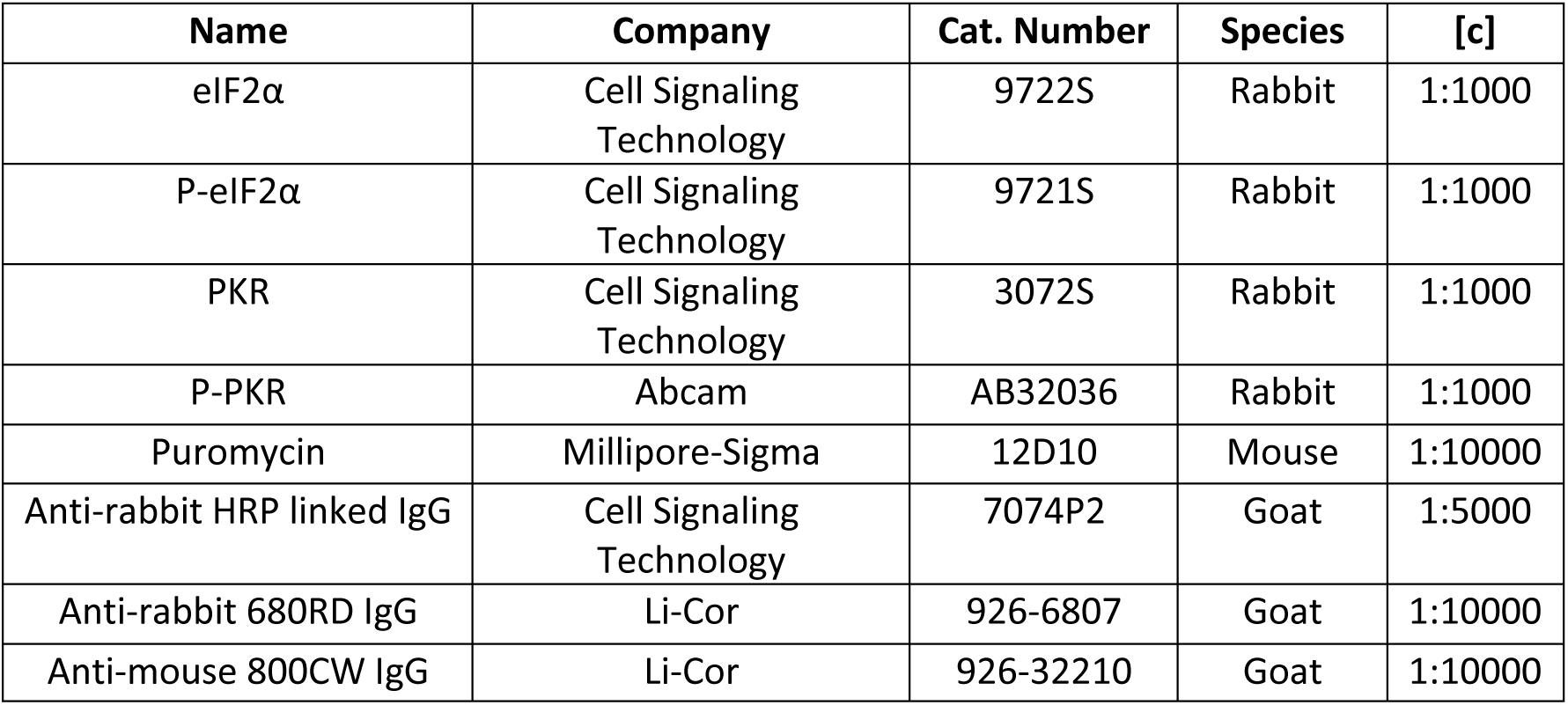

| Name | Company | Cat. Number | Species | [c] |
| --- | --- | --- | --- | --- |
| eIF2α | Cell Signaling Technology | 9722S | Rabbit | 1:1000 |
| P-eIF2α | Cell Signaling Technology | 9721S | Rabbit | 1:1000 |
| PKR | Cell Signaling Technology | 3072S | Rabbit | 1:1000 |
| P-PKR | Abcam | AB32036 | Rabbit | 1:1000 |
| Puromycin | Millipore-Sigma | 12D10 | Mouse | 1:10000 |
| Anti-rabbit HRP linked IgG | Cell Signaling Technology | 7074P2 | Goat | 1:5000 |
| Anti-rabbit 680RD IgG | Li-Cor | 926-6807 | Goat | 1:10000 |
| Anti-mouse 800CW IgG | Li-Cor | 926-32210 | Goat | 1:10000 |

## Puromycilation of nascent peptides

Cells were grown in culture medium until 80% confluency in 6-well plate for a day (one well per condition). Cells were then treated with activating blue light illumination accordingly until 30 min prior to the end of their treatment, at which point, the culture media was replaced with media containing 18 µM puromycin (Gibco, A1113803) and the cells were incubated for an additional 30 min before sample collection and analysis as described in the Western blotting section.

### Immunofluorescence

Cells were grown on fibronectin (Sigma, F1141) coated glass bottom 96 well plates (Cellvis, P96-1.5H-N), washed with PBS, and fixed using 4% paraformaldehyde (PFA, Fisher Scientific, AA433689L) in PBS for 10 minutes at room temperature under dark conditions. Fixed cells were washed with PBS and permeabilized with blocking buffer (0.05% saponin, 0.5% BSA, 50 mM, NH_4_Cl in PBS) for 30 min at room temperature. The fixed cells were incubated with primary antibodies diluted in blocking buffer overnight at 4°C. The next day, the cells were washed gently 3x with PBS-T, followed by secondary antibody incubation for 1 hour at room temperature in dark conditions, followed by another round of 3x PBS-T washes. The prepared fixed cells were imaged as detailed in Microscopy imaging.

List of antibodies used in immunofluorescence analyses:

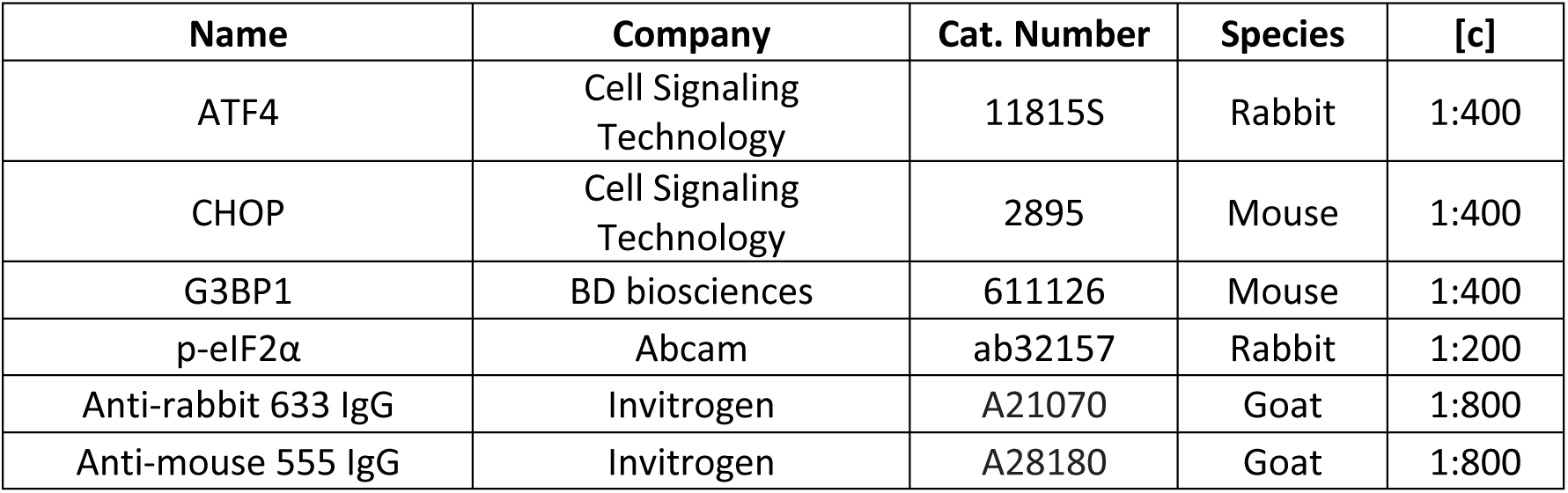

| Name | Company | Cat. Number | Species | [c] |
| --- | --- | --- | --- | --- |
| ATF4 | Cell Signaling Technology | 11815S | Rabbit | 1:400 |
| CHOP | Cell Signaling Technology | 2895 | Mouse | 1:400 |
| G3BP1 | BD biosciences | 611126 | Mouse | 1:400 |
| p-eIF2α | Abcam | ab32157 | Rabbit | 1:200 |
| Anti-rabbit 633 IgG | Invitrogen | A21070 | Goat | 1:800 |
| Anti-mouse 555 IgG | Invitrogen | A28180 | Goat | 1:800 |

### Microscopy imaging

All microscopy imaging, live or fixed, was done using a W2 SoRa spinning-disk confocal microscope equipped with an electron-multiplying charge-coupled device camera (Andor, DU-888) and an incubation chamber maintaining cells at 37°C and 5% CO2. All cells that were imaged were plated on fibronectin (Sigma, F1141) coated glass bottom 96 well plates (Cellvis, P96-1.5H-N). Images were captured using a 40x NA 1.25 water immersion objective and a 60x NA 1.2 water immersion objective.

### Optogenetic stimulation

Engineered photo-switchable proteins were activated using 455 nm blue light. For Western blot experiments with maximal light activation of the pathway, we utilized a custom 36 LED plate mounted on a box lined with aluminum foil, allowing for equal light distribution throughout the box. For confocal imaging, light was delivered through a 488 nm laser at 100% power using a 40x NA 1.25 water immersion objective. For spatial patterning of light, we utilized purpose-built microscope-mounted LED-coupled digital micromirror devices (DMDs) triggered via Nikon NIS Elements software, allowing us to specify the exact parameters of light delivery at each coordinate of our imaging plane. For precisely modulated IF experiments, we utilized the benchtop LED array light delivery device (optoPlate), per authors instructions (Bugaj and Lim, 2019). The light delivery parameters for the optoPlate were purpose-set for each experiment, where stimulating light was continuously ON or OFF, without pulsing, and 0.2-7.5 mcd range of intensities applied translates as 3-96 on the intensity setting of the optoPlate software. The light illumination sequence of each well included an OFF (dark) period at the start, so that all well illuminations would end at the same time.

### Image analysis

All quantification of images followed the same general flow from creating a TIFF library of images to clean-up and analysis using custom code on MATLAB (Supp. Fig. 3C). Nikon-generated ND2 files were channel separated and converted to TIFF files using ImageJ FIJI macro (ImageJ). The TIFF library of images was organized into separate stacks based on applied treatment (i.e. different intensity of activation, etc.) over time on MATLAB, for ease of access and indexing. All images were then cleaned up to reduce noise through background subtraction and deletion of super bright pixels. DAPI stained nuclei were segmented to identify nuclei as follows: First, we Gaussian-blurred the images, then used threshold detection to identify edges, and watershed transformed the nuclear mask in case of connected nuclei. The nuclear masks were quality checked and the threshold was adjusted manually in case of low quality masking. For nuclear localized information, we applied the nuclear mask to the channel of interest and collected information of every nucleus separately. For cytoplasmic information, we dilated the nuclear mask and removed the mask itself from the dilated mask to quantify the ring around the nucleus as cytoplasmic information. For condensate information, we identified clusters through both “Laplacian of Gaussian” edge detection and intensity threshold detection and combined the information to create a condensate mask. We also removed non-condensate information from our mask as follows: outside our size threshold and nuclear localized. Mean intensity, location and area of all objects were saved for quantitative analysis on a single cell level for nuclear and cytoplasmic fluorescence normalized to no treatment control. Condensate data was quantified by summing total condensate intensity across an imaging field and normalized to the number of nuclei detected within said image field to obtain the mean condensate fluorescence per cell.

### Bulk RNAseq

#### Sample treatment and extraction

H4 cells carrying Cry2Olig-PKR cells were plated on fibronectin treated 96-well glass-bottom plate. Cells were illuminated according to experimental set-up using the optoPlate device set to continuous illumination intensity of 7.5 mcd as mentioned in Optogenetic stimulation. Cells were immediately lysed using 120 µL Trizol (Gibco, 15596026) per well, under red light, and incubation for lysis for 5 min at room temperature. RNA was extracted according to the manufacturer’s protocol.

#### Illumina library preparations

Bulk RNAseq Illumina libraries were prepared following an adaptation of the single-cell method “CEL-Seq2” where we treated every illumination duration as a single cell (Hashimshony et al., 2016; Wangsanuwat et al., 2020). Briefly, sample RNA was reverse transcribed with Cel-Seq2 primer #4 (25 ng) using Superscript II reverse transcriptase. Second Strand Synthesis (SSS) was carried out via *E. coli* ligase (Invitrogen, Cat. # 18052–019) and DNA polymerase I (Invitrogen, Cat. # 18010–025) in SS buffer for 2 hours at 16°C. Next, cDNA was purified with 1:1 AMPure XP DNA beads (Beckman Coulter, Cat. # A63881), and washed with 80% ethanol twice, before elution into nuclease-free water and vacufuge concentrated. *In vitro* transcription was then carried out using MEGAscript T7 kit (Thermo Fisher Scientific, AMB13345) for 13 hours at 37°C, followed by RNA treatment using EXO-SAP (Thermo Fisher Scientific, 78200.200.UL) for 15 min at 37°C, and finally the samples were fragmented (200 mM Tris-acetate (pH 8.1), 500 mM KOAc, 150 mM MgOAc) at 94°C for 3 min before being quenched by 0.5 M EDTA. The fragmented RNA was size selected with 0.8x AMPure RNA beads (RNAClean XP Kit, Beckman Coulter, Cat. # A63987) and eluted in nuclease-free water.

#### Bioinformatic analysis

Paired-end sequencing of the libraries was performed on an Illumina NextSeq 500, and hits were quantified and z-score normalized per gene. Transcriptomic data was then analyzed through custom pipeline on python packages “numpy” and “pandas”. As data preparation, all genes that were missing entry at any time were removed from data matrix. We then applied SVD function from “numpy” to gene matrix, where rows were distinct genes and columns were duration of stimulation in increasing order, then sorted the data based on U matrix of SVD decomposition. We also applied centroid clustering on the gene matrix through clustermap function of “seaborn” package.

### Simulating the mathematical model of ISR

Model simulations were conducted in Python 3 using the Anaconda 4.1.3 using the packages “numpy”, “pandas” and “seaborn”. Numerical simulations were conducted using the “odeint” function from the SciPy v1.8.0 scipy.integrate module. All computational experiments can be found in Supplementary Data (Data S1).

Upon defining the ISR model as ISR_linear and ISR_U functions with all the state variables and parameters listed in Fig. S4A-D, for linear and cooperative models, respectively. We defined non-linear variable light modulation using Ustep(t), as a range of [0] to [1] as a proxy for illumination at specific intensities. PKR and p-PKR defined in our model refers to our engineered opto-PKR and its phosphorylated form, and Ustep(t) was incorporated into these state variables. Initial conditions of the state variables were set to 0 for phosphorylated proteins (p-PKR, p-eIF2α) and GADD34, while non-phosphorylated proteins (PKR, eIF2α) were set to 1, as shown at time zero of simulated step response (Fig. 4D). All state variables were then plotted using matplotlib plotting library and/or seaborn.heatmap library.

For step response simulation, we set Ustep to [1] at time 2 hrs (of modeled time), and let the simulation run through 20 hours. For intensity scan simulation, we queried the magnitude of Ustep from [0.03] to [1], by increasing it from [0] at 2 hrs of modeled time. For the input-recovery ratio simulation, we pulsed Ustep to [1] starting at 2 hours for corresponding length of modeled time (1.5, 3, or 6) hrs, before pulsing for 30 min (0.5 hrs) at time 10 hours. For simulated stress memory landscape, we log sampled light durations (Ustep == 1) and recovery durations (Ustep == 0), from [0.5] to [10] hours and [2] to [48] hours, respectively, through nested for loop solving the ISR model with odeint for each condition.

### Gene ontology analysis

Genes of interest were obtained from the U vector of SVD analysis. The top 500 genes were analyzed for over/under representation in biological processes of *Homo* Sapiens using GO Enrichment Analysis tool (Panther, “geneontology.org’) (Ashburner et al., 2000; Gene Ontology Consortium, 2021; Mi et al., 2019). The dataset used for this analysis was GO biological processes complete set and performed Fisher’s exact test with correction for false discovery rate. All biological processes found significant were collapsed onto its parent families.

---

**Supplementary Figure 1.**
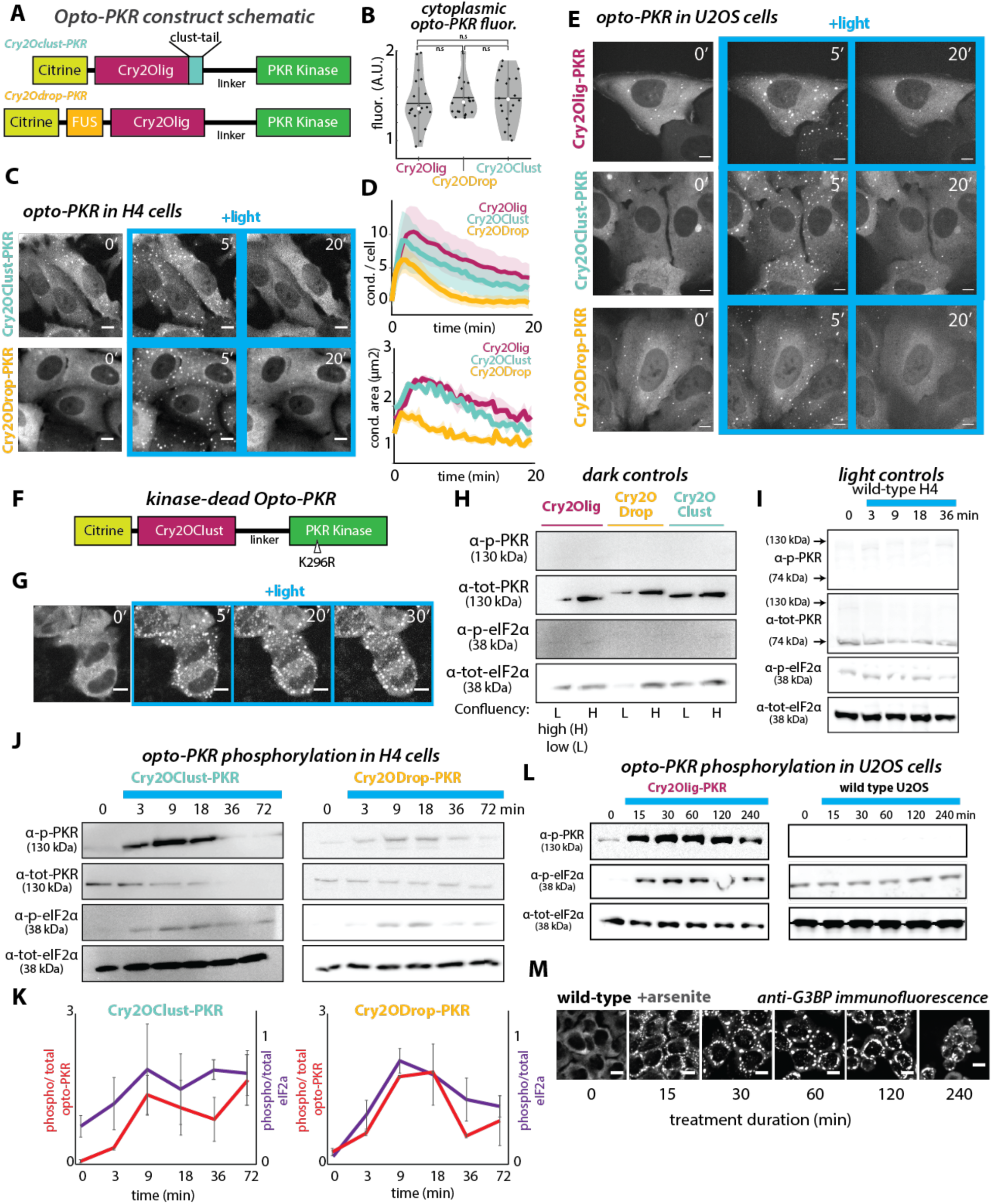
(A) Construct design and architecture of opto-PKR variants. Modifications added to the Cry2 domain are clust-tail or FUS domains. Fused to human PKR at residue 170 to 551. (B) Violin plot of cytoplasmic opto-PKR fluorescence levels quantified from live cell imaging. Each black dot indicates a sample, the horizontal line indicates the mean, the white dot indicates the median. All cell lines had non-significant variation in student’s t-test, p > 0.05. n = 20. (C) Live cell imaging of opto-PKR variants in H4 cells activated under constant 448 nm illumination. Images are the maximum intensity projection of 5 Z-planes taken through the cell. Scale bar = 10 µm (D) Quantification of live cell imaging (Fig. 1C, Fig. S1C), in terms of opto-PKR condensate characteristics of number per cell (top) and size (bottom) over time. (E) Live cell imaging of opto-PKR variants in U2OS cells activated under constant 448 nm illumination. Images are the maximum intensity projection of 5 Z-planes taken through the cell. Scale bar = 10 µm. (F) Construct design and architecture of kinase-deficient opto-PKR (opto-PKR-KK, harboring a K296R mutation in its kinase domain). (G) Live cell imaging of H4-opto-PKR-KK cell lines under continuous 445 nm laser activation, same as in C. Images are the maximum intensity projection of 5 Z-planes taken through the cell. Scale bar: 10 µm. (H) Western blot of H4-opto-PKR and variants in dark conditions. (I) Western blot of H4-WT cells under continuous blue light stimulation. (J) Representative Western blots of phosphorylated and total opto-PKR and eIF2α in cells bearing opto-PKR variants under continuous blue light stimulation. Light treatment over 72 mins. (K) Quantification of the data from J. Phosphorylation signal was normalized as ratio to the total protein (PKR or eIF2α) at each time point. N = 3. Error bars: SEM (L) Western blots of phosphorylated and total opto-PKR and eIF2α in U2OS-opto-PKR and U2OS-WT cells under continuous blue light stimulation. Light treatment over 240 mins. (M) Immunofluorescence analysis of the SG marker G3BP1. Representative images of H4-WT cells exposed to sodium arsenite in the dark (500 µM). Scale bar = 10 µm.

**Supplementary Figure 2.**
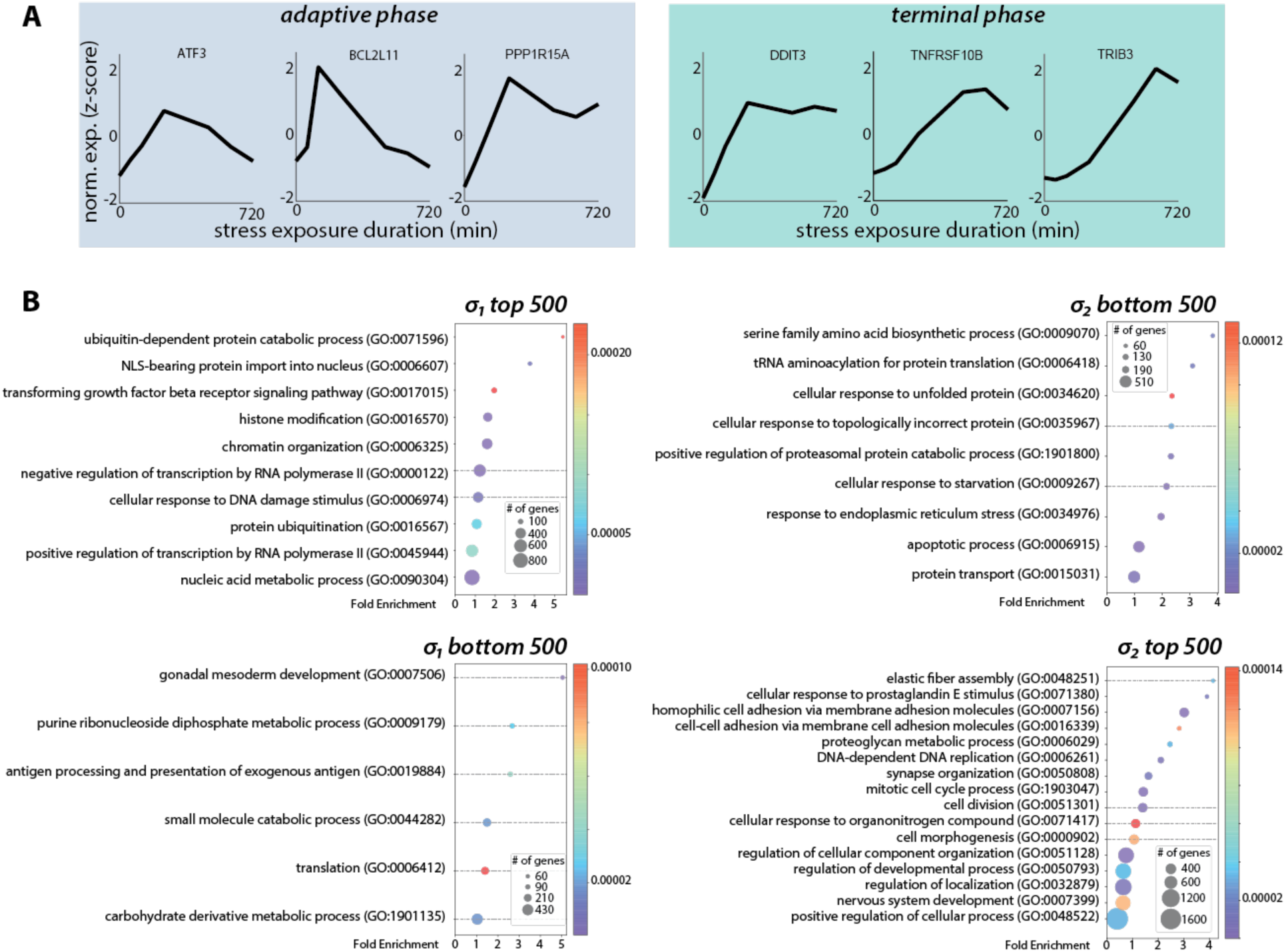
(A) Gene expression dynamics of single genes of interest from RNAseq hits. (B) Gene ontology enrichment analysis of the 500 genes shown in Fig. 4C, showing functional processes of the genes that contributed most to the dynamical transcriptional modes observed in the ISR.

**Supplementary Figure 3.**
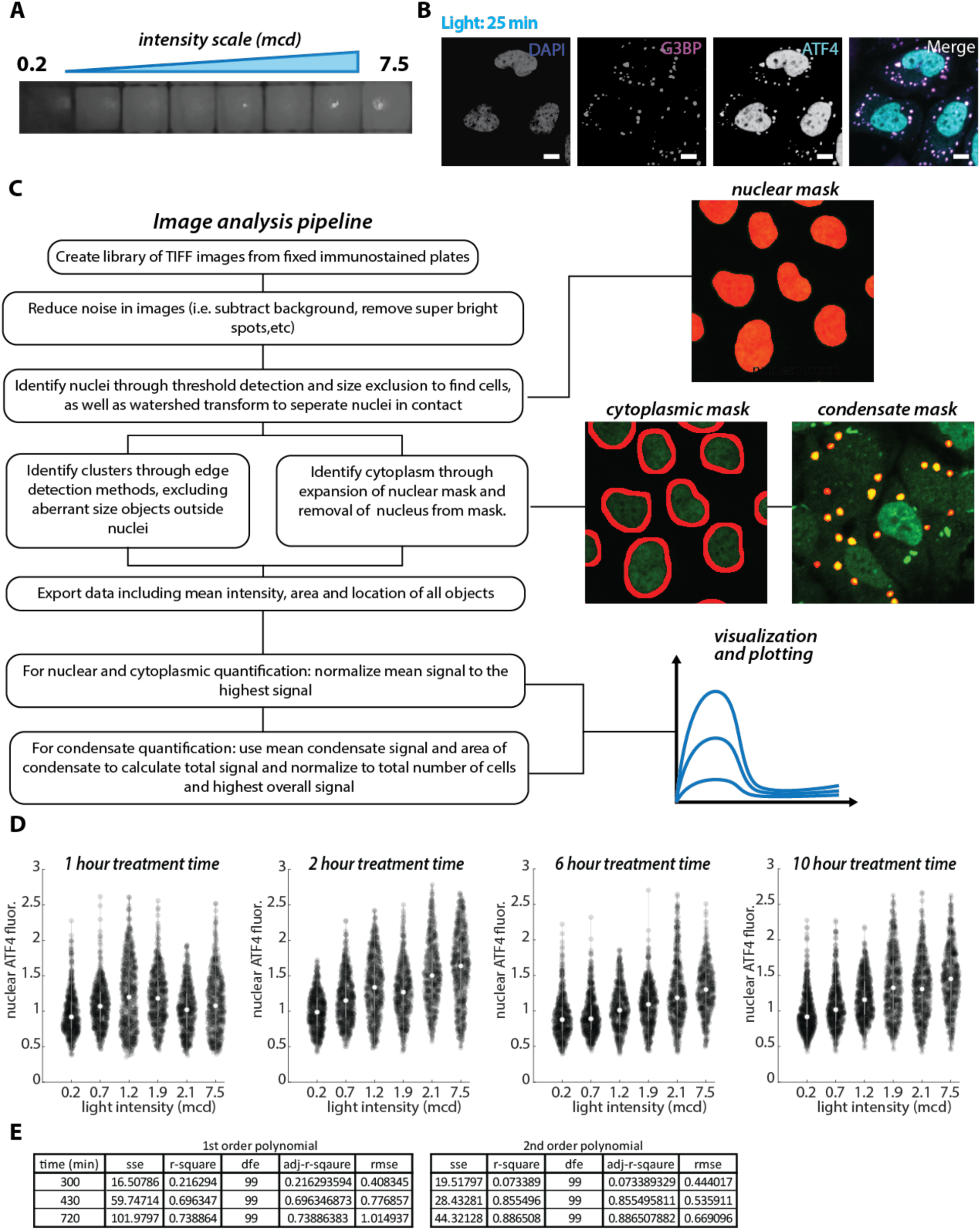
(A) Image of the optoPlate device at relevant light intensities ranging from 0.2 mcd to 7.5 mcd. (B) Representative immunofluorescence images of H4-opto-PKR cells under blue light stimulation at 3.8 mcd for 25 min. Stained for nucleus (DAPI), G3BP1 and ATF4. Scale bar = 10 μm. (C) Flow diagram of actions in our automated image analysis pipeline. In example images, masking is shown in the red channel and fluorescence data in green. (D) Violin plots of nuclear ATF4 fluorescence under continuous exposure at varying intensities over time (as illustrated in Fig. 3B). Nuclear ATF4 fluorescence normalized to no light illumination control. Each dark dot represents quantified nucleus, and the mean is indicated by a white dot. (E) Table of goodness-of-fit statistics for both first (f(x) = ax+1) and second order (f(x) = ax^2+1) polynomial fits shown in Fig 3J.

**Supplementary Figure 4.**
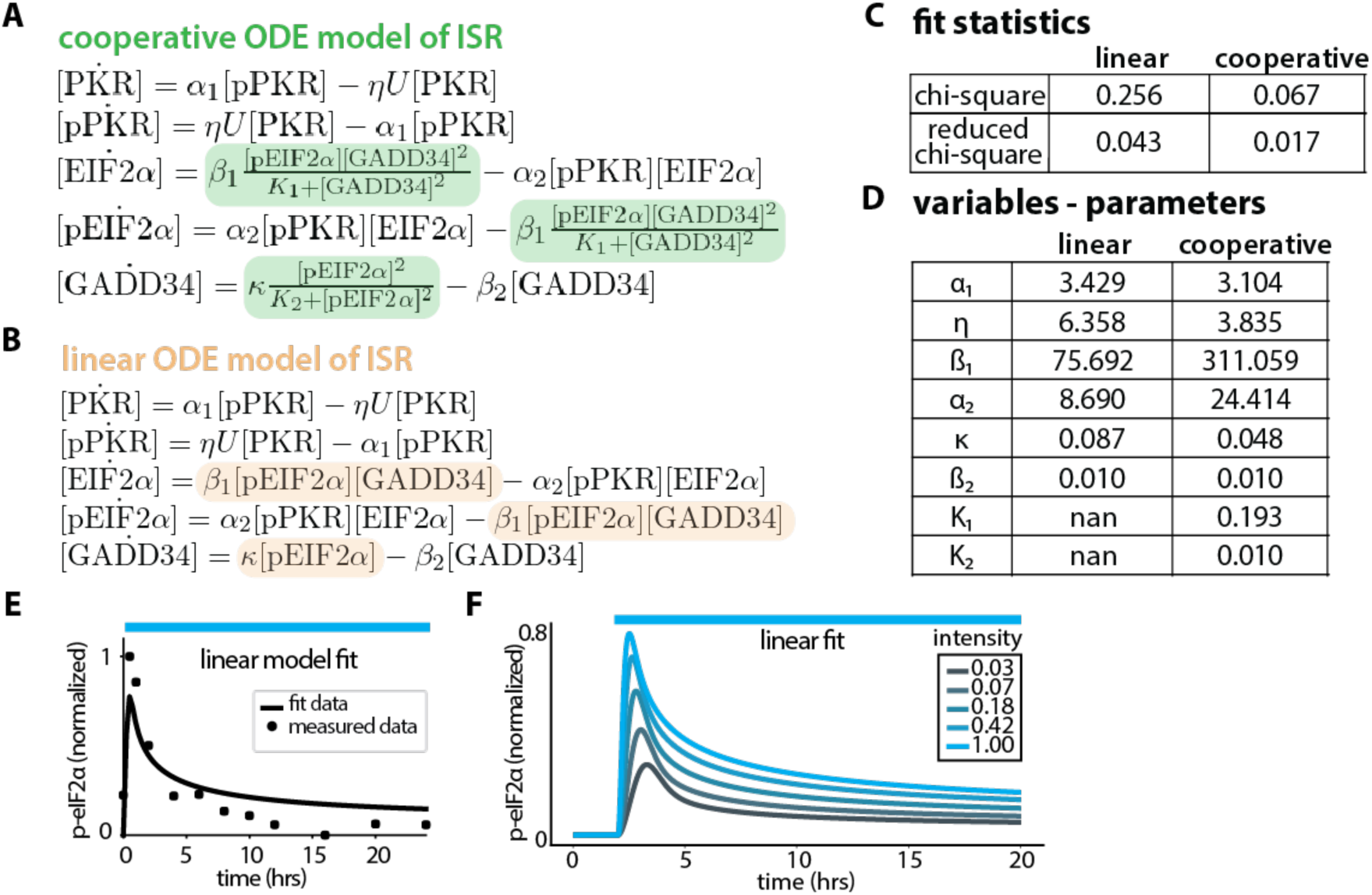
(A,B) Sets of ordinary differential equations (ODE) defining interactions of state variables shown in Fig. 4A. Variable “U” represents light illumination. Cooperative (A) and linear (B) elements are highlighted. (C) Goodness-of-fit statistics for the linear and cooperative models to experimental data as shown in Fig 4B and Supp Fig. 4E. (D) Table of parameter values for best fit of linear and cooperative models to experimental data. (E) Best fit of simulated data through linear ODE model to experimental data for p-eIF2α, where line is simulated data and points are experimentally measured data. (F) Simulated data through linear ODE model showing changes in p-eIF2α levels in response to decreasing illumination intensity, adjusted as part of the differential equation representing opto-PKR.

**Supplementary Figure 5.**
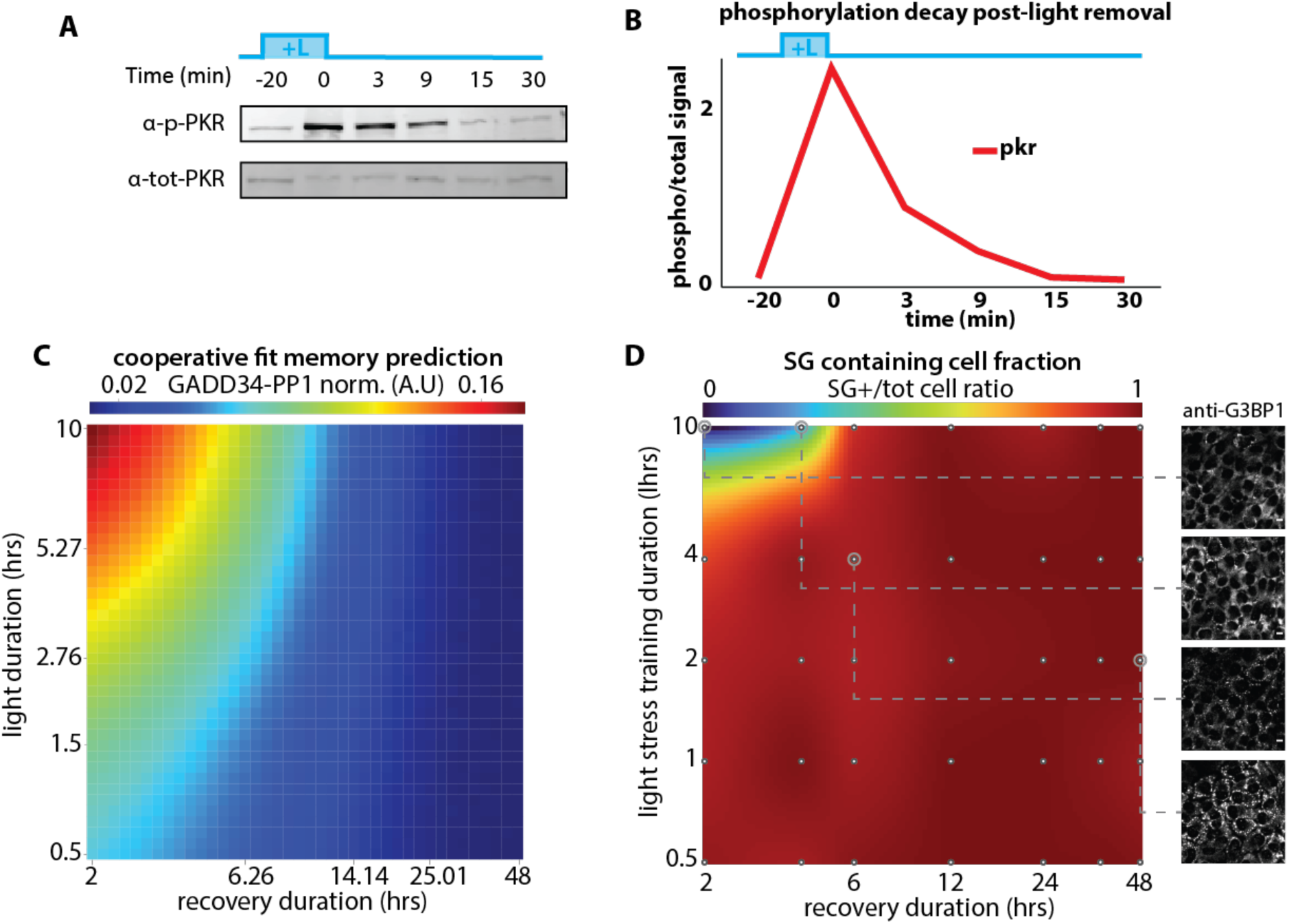
(A) Western blot of opto-PKR under no light treatment post-illumination to measure phosphorylation decay. Pre-treated H4-opto-PKR cells with 20 min of blue light, before following opto-PKR levels over time. Time -20 min = no blue light exposure and time 0 min = post 20 min of blue light exposure and start of decay measurement. (B) Quantification of Western blot in C. Phosphorylation signal normalized as fold change in comparison to the total protein at each time point. (C) Heatmap of simulated stress memory landscape in terms of the GADD34 response to the 10 min challenge. Total duration of conditioning varies. (D) Interpolated heatmap of the fraction of cells that formed SGs in response to challenge after conditioning. Experimental data quantified from immunofluorescence staining for G3BP1 and ratios of cells with SGs compared to total cell count. Each dot represents a sample point. Interpolation was done using a modified Akima cubic Hermite method. Insets are representative immunofluorescence images at each grey circled sample point.

## Notes

### Competing Interest Statement

The authors have declared no competing interest.

